# Glioma cell migration in confined microchannels via a motor-clutch mechanism

**DOI:** 10.1101/500843

**Authors:** Louis S. Prahl, Maria R. Stanslaski, Pablo Vargas, Matthieu Piel, David J. Odde

## Abstract

Glioma tumor dispersion involves invading cells escaping the tumor bulk and migrating into the healthy brain parenchyma. Here, they encounter linearly aligned track-like tissue structures such as axon bundles and the perivascular space. These environments also contain micrometer-scale pores that impose mechanical confinement on invading cells. To study glioma cell migration in an *in vitro* system that reproduces some of these features, we used microfluidic devices with 60 μm^2^ cross-sectional area channels that confine cells into one-dimensional (1D) tracks. Individual cell tracking revealed strongly persistent migration at a mean rate of 8.5 ± 0.33 nm s^-1^. Notably, a 1D computational cell migration simulator predicts migration behaviors of glioma cells without significant adjustment of parameters estimated from previous experiments on two-dimensional (2D) substrates. Pharmacological inhibitors of integrin-mediated adhesions, myosin II activation, or drugs targeting F-actin assembly or microtubule dynamics influence migration consistent with simulations where relevant parameters are changed. These results suggest that cell parameters calibrated to a motor-clutch model on 2D substrates effectively predict 1D confined migration behaviors *a priori*. Our results outline a method for testing biophysical mechanisms of tumor cell migration in confined spaces and predicting the effects of anti-motility therapy.

## Introduction

Tumor cell dispersion into healthy brain tissue promotes the rapid disease progression and recurrence of glioblastoma (GBM, grade IV glioma) (1, 2). Cells invading the brain parenchyma encounter a dense extracellular matrix (ECM) rich in hyaluronic acid and containing densely packed cells (3, 4). Specific invasion routes include aligned structures including axon bundles, astrocyte processes, and blood vessels (4, 5) that guide cell migration along linear tracks. Perivascular spaces and spaces along the glia limitans contain rigid basement membranes rich in laminin and collagen (5). Many of these environments require invading GBM cells to squeeze through constricted channels or micrometer-scale pores between other cells (6). More generally, interstitial spaces within or between tissues present similar mechanical challenges to invading tumor cells (5), as well as immune cells performing tissue surveillance (7).

Traditional two-dimensional (2D) *in vitro* cell culture assays lack aligned structures and confined pores found in tissue. Architectural features can guide cell migration; for example, cells moving with aligned guidance cues infrequently reverse direction (8–11). One-dimensional (1D) topographical cues of similar dimension can be replicated on grooved silicon wafers (12), suspended polystyrene fibers (13), aligned collagen gels (14), or using micro-contact printing of ECM proteins (15, 16). However, most of these culture systems still do not enforce the significant degree of confinement encountered by cells migrating *in vivo*. To overcome these limits, photolithography and polydimethylsiloxane (PDMS) replica molding can create reproducible micrometer-scale channels (8–11, 17–21) enabling mechanistic cell migration studies in a reproducible fashion.

The classical mechanism of cell migration involves cyclic steps of F-actin based protrusion, adhesion, myosin II-based contraction, and de-adhesion that permits translocation of the cell body (22). Physics-based models, such as the motor-clutch model (23), combine these into a computational framework that predicts cell force generation as a function of myosin II motor activity and binding kinetics of molecular adhesion receptors that recognize ECM ligands. A cell migration simulator (CMS) based on the motor-clutch model was shown to predict GBM cell migration behaviors in environments with varying chemical and mechanical properties (13, 24, 25) as well as the effects of drugs targeting cellular components involved in motility (25, 26). Given that GBM migration speed correlates inversely with patient survival (24), CMS predictions could inform *in silico* estimates of tumor progression and response to therapy. However, the CMS has not been rigorously tested on data generated in the mechanical confinement conditions cells encounter *in vivo*. Confinement can variably influence cell migration, including triggering fast, polarized migration in low adhesion conditions (27) or impeding migration when cells must deform their nuclei to pass through micron-scale constrictions (28). Recent works also question whether confined tumor cells use actin-myosin contractile machinery and integrin-mediated adhesions (22) or osmotic pressure (29, 30) as their primary means of force generation.

Typical methods of inhibiting migration on 2D substrates may have no effect on cells in confinement (11, 29), which highlights the need to mechanistically evaluate these treatments in a tissue-appropriate context. Microtubules are involved in tumor cell migration and proliferation – processes that drive tumor dispersal – and are frequently targeted in chemotherapy regimens by microtubule-targeting agents (MTAs). MTAs potently inhibit glioma cell migration on 2D substrates (26, 31–33) and slow migration of some cells in confinement (8, 11), but not all cell types appear to be sensitive (19). Microtubules regulate a number of key signaling pathways involved in migration (34), but whose roles in confined migration are controversial, including pathways that activate myosin II (35) and promote integrin-mediated adhesion (36). Systematic, modeling-based approaches could be used to parse out the effects of MTAs and connect them to the appropriate biophysical parameters.

To develop a modeling approach capable of systematically testing both the effects of environmental structures and anti-motility therapy, we made simple modifications to an existing CMS (24, 25) that reflect the geometric limits of a confined microchannel. We used PDMS microchannel devices to create experimental conditions of mechanical confinement and found that the human U251 GBM cell line spontaneously migrates persistently along the channel axis, consistent with simulation predictions. Pharmacological inhibitors of myosin II and integrin-mediated adhesion also influence migration speed, consistent with biophysical modeling predictions. Next, we demonstrate that MTAs and inhibitors of actin assembly effectively slow migration, consistent with altered protrusion dynamics parameters in simulations. Our results imply that parameter sets derived from experimental measurements on 2D substrates (25, 26) are valid for GBM cells in confined environments, and that therapies targeting the motor-clutch system or MTAs are viable ways to slow GBM cell migration in tissue-like confinement.

## Materials and Methods

### 1D cell migration simulator

Previous implementations of the cell migration simulator (CMS) either considered a cell migrating on a 2D unconfined surface with varying adhesiveness or mechanics (24–26), or explicitly modeled the geometric constraints of suspended fiber networks (13). In the present study, we modified the CMS to model cell migration in a single spatial dimension, hereafter referred to as the 1D CMS (a schematic is shown in **Figure 1A**). As with previous versions, the 1D CMS contains j protrusion modules, each functioning as an instance of the motor-clutch model (23). Modules contain n_clutch,j_ adhesion clutch molecules, each of which is modeled as a Hookean spring with stiffness κ_clutch_. Clutches bind to a compliant substrate at a reference point x_ref,j_. The substrate is also modeled as a linear (Hookean) spring with stiffness κ_sub_. Clutches bind at a first-order rate k_on_. Unbinding of the i^th^ clutch in a particular ensemble occurs at a rate k_off,i_ that scales with force by a single exponential (37), also known as a slip bond.

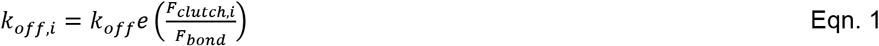

**Figure 1.**
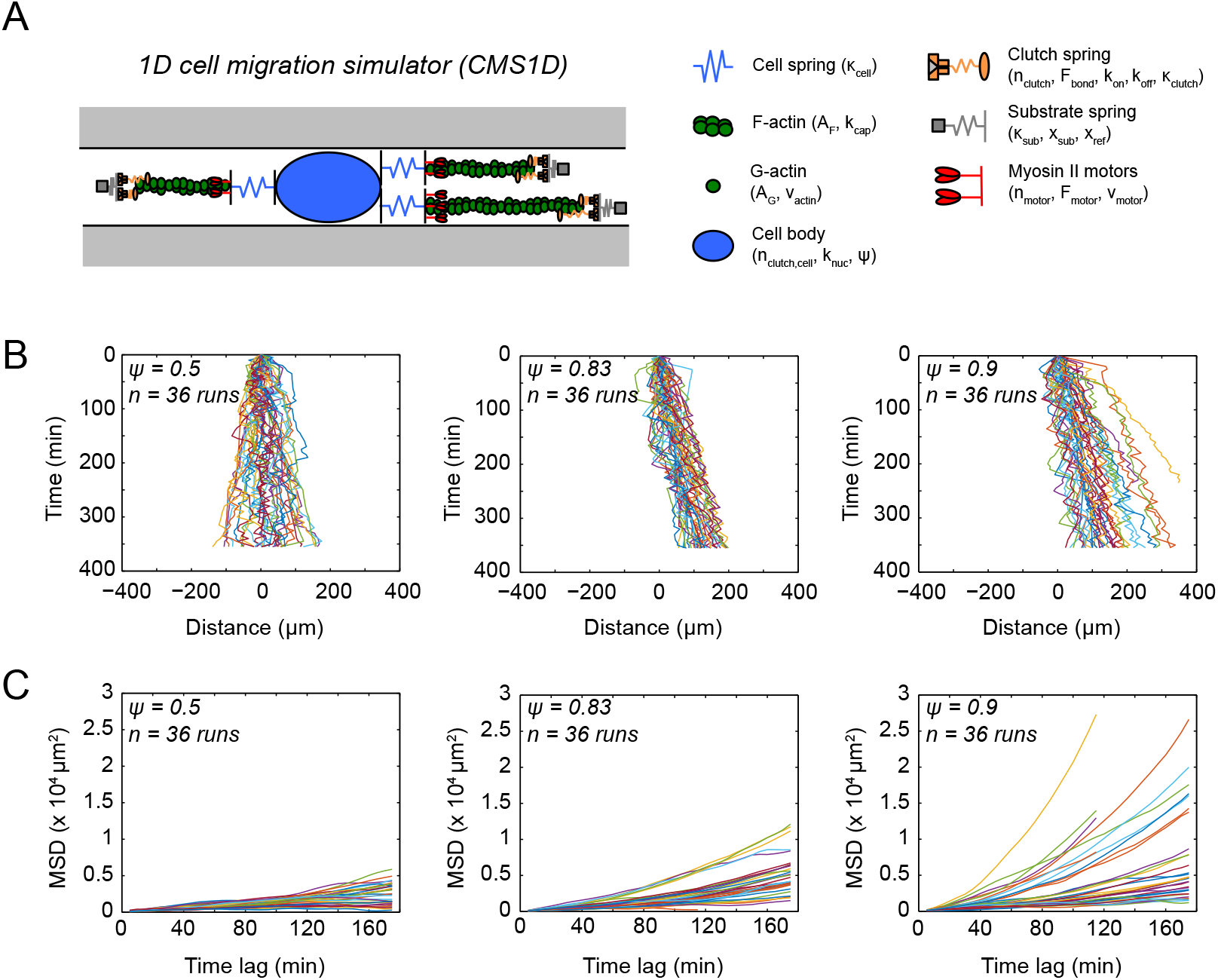
Description and migration dynamics of a 1D cell migration simulator. (A) Schematic of a 1D cell migration simulator (1D CMS) within a confined channel (denoted by gray walls). Modules containing motors (red), clutches (orange), and F-actin (green) attach to a central cell body (blue oval) by compliant springs (blue). Modules attach to and extend a compliant substrate spring (grey) at their distal end. Cell body clutches (not pictured) associate with the cell center, x_cell_. An actin mass balance governs module nucleation with rate constant (k_nuc,0_) and maximum actin polymerization speed (v_actin,max_) by previously described relationships (25). Module capping (k_cap_) terminates polymerization and facilitates module shortening and turnover and ψ governs the probability of new protrusions being generated in the +x direction. **Supporting Material** contains a detailed model description. (B) Simulation position as a function of time for individual 1D CMS runs where ψ = 0.9, 0.83, or 0.5, n = 36 simulated trajectories are shown for each condition. Initial position is marked at x(τ = 0) = 0. All other simulation parameters are kept constant between conditions and reported in **Table S1**. (C) MSD versus time lag for the 1D CMS trajectories in panel B.

In Eqn. 1, k_off_ is the unloaded unbinding rate, F_bond_ is the characteristic bond force, and F_clutch,i_ is the force on the i^th^ clutch. Modules also contain n_motor,j_ myosin II motors, each of which is capable of generating F_motor_ stall force. Motors slide an F-actin bundle to generate retrograde flow (v_flow_), which extends clutches that are bound to the substrate. As forces build on the substrate through bound clutches, F-actin flow slows from the myosin II motor unloaded velocity v_motor_ by a linear force-velocity relationship (23).

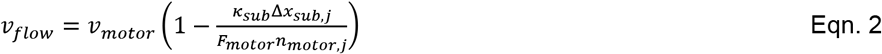

In Eqn. 2, Δ_xsub,j_ is the substrate spring displacement on the j^th^ module. Each module contains a rigid bar of F-actin with length A_F,j_. Modules extend by actin polymerization (v_actin_) from a soluble pool of G-actin subunits, A_G_. Actin polymerization (v_actin_) is defined by a maximum actin polymerization velocity v_actin,max_, A_G_, and the total available actin length in the cell, A_total_.

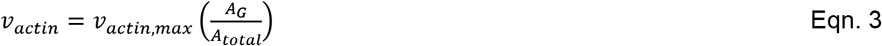

A mass balance constrains A_total_ and A_G_ in a simulated cell with n_module_ modules.

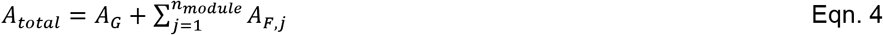

F-actin is depolymerized when it passes the motor position within modules (x_motor,j_) and returns to the G-actin pool in the cell. Module clutches also unbind when their bound position (x_clutch,i_) extends past x_motor,j_. Modules contain a compliant cell spring with stiffness κ_cell_ that represents the nucleo-cytoskeletal compliance of the cell and connects the module motors to the central cell body. A force balance applies to each module spring, clutches, and cell spring.

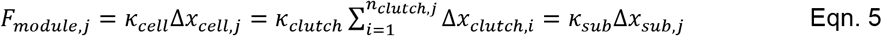

Within the cell body, an ensemble of n_clutch,cell_ clutches transmit force to the substrate under the cell body in the same way as module clutches. F_cell_ represents the force from extension of the cell substrate spring Δ_xsub,cell_

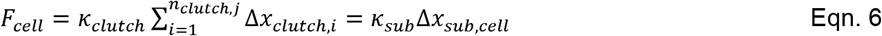

Cell body clutches are not subject to direct myosin II forces as module clutches are, but extend as the cell position x_cell_ is updated by force balance on modules and the cell body.

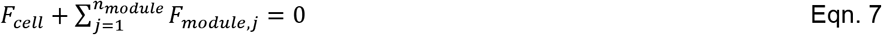

Module capping occurs at a first-order rate k_cap_ and capped modules no longer extend by actin polymerization. Nucleation of new modules occurs at a rate k_nuc_, and k_nuc_ is a function of the basal nucleation rate (k_nuc,0_) and the available G-actin pool raised to the fourth power (38).

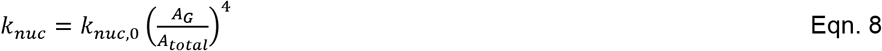

New modules have initial length L_cell_ and are assigned motors and clutches as a fraction of the available pool of each within the cell (25). Modules that shorten past a minimum length L_min_ are eliminated and their contents (motors, clutches, F-actin) are returned to the common cellular pool.

In the previous CMS, modules are randomly nucleated at a uniformly distributed angle θ within the 2D plane. Module nucleation in 1D can only occur in the +x or –x direction, so we defined a cytoskeletal polarity factor ψ that represents the probability that a new module will be generated in the +x direction. The complementary probability 1-ψ is the probability that a new module is generated in the –x direction. The case where ψ = 0.5 represents an equal likelihood of module generation in the +x and –x directions, which can be thought of as a “non-polarized” cell. In a cell where ψ < 0.5, the polarity would bias module nucleation in the –x direction. Essentially, the probability of module nucleation in the +x direction follows a Bernoulli distribution with parameter ψ. All simulation parameter values are given in **Table S1** in the **Supporting Material** and specific changes are described in figure legends.

### Simulation implementation and analysis

In simulations, event selection was determined using a Gillespie stochastic simulation algorithm (SSA) (39) with a unique, randomly generated seed for each run. The order of events in each step of the model algorithm was as previously described (25). Simulation initial conditions included two modules oriented in the +x direction and one module oriented in the –x direction (each module of length L_cell_) and all clutches unbound. Individual runs represent data collected over 5-7 hours of simulated time. Simulations were run in Matlab (The MathWorks Inc., Natick, MA) on in-house computing cores or using high performance computing cores in the Minnesota Supercomputing Institute.

To quantify simulated cell migration, cell body position (x_cell_) was sampled every 5 minutes of simulated time. We excluded the first hour of simulated data from analysis and subsequent data were filtered to remove large displacements in cell position that result in velocities >1 μm s^-1^ (25). MSD was calculated from cell body position using the overlap method (25, 40) and fit to variants of the random walk model to extract parameters related to cell migration (described in the main text) using a custom Matlab analysis script. The first half of MSD versus time lag data was used for each individual cell fit.

### Microchannel master molds

Microchannel devices were designed using computer-aided design software (AutoCAD; Autodesk, San Rafael, CA). Channel architectures were based on previously described designs (41, 42). Devices feature three rectangular seeding ports, each marked with 15 μm diameter pillars evenly spaced at 25 μm intervals and with opposite rows of channels aligned towards the device exterior. Entrances to channels are 100 μm long funnels that taper from an initial width of 18 μm to a final channel width of 12 μm. Chromium-plated quartz photomasks were generated based on these designs by the Minnesota Nano Center (MNC) facilities. Device molds were fabricated on 4” silicon wafers using epoxy-based SU-8 2005 negative photoresist (MicroChem Corp., Wesborough, MA) and soft photolithography. A 5 μm thick photoresist layer was spin coated on wafers using a CEE 100 spin coater (Brewer Science, Rolla, MO), followed by a 2 minute soft bake at 95°C. Wafer surfaces were exposed to UV radiation at 12 mJ s cm^-2^ for 10 seconds using a MA6 mask aligner (Karl Süss MicroTec, Garching, Germany). Following a 3 minute post-exposure bake at 95°C, excess photoresist was removed using SU-8 developer (MicroChem). After curing, the height of individual photoresist structures was verified using a P-16 stylus profilometer (KLA-Tencor, Milpitas, CA). To minimize PDMS adhesion and facilitate device removal, wafers were treated with trichloro(1H,1H,2H,2H-perfluorooctyl)silane (Sigma, St. Louis, MO) under vacuum before casting.

### Device fabrication and assembly

Sylgard^®^ 184 PDMS elastomer (Dow Corning, Midland, MI) was mixed in a 10:1 (base:curing agent) ratio, poured on wafers or epoxy replicas and allowed to cure at 75°C for 2 hours. Devices were cut and peeled from molds, seeding ports were added using a 3 mm radius circular punch (Ted Pella Inc., Redding, CA), and devices were manually resized to a 1.5 x 1.0 cm footprint centered around the seeding ports using a razor blade. Cut and resized devices were cleaned with adhesive tape and 70% ethanol in a sonic bath for 1 minute and then allowed to dry under a stream of compressed air. Devices and 35 mm glass bottom dishes (20 mm No. 0 coverglass, MatTek Corp., Waltham, MA) were activated with a plasma cleaner (PDC-32 G, Harrick, Ithaca, NY) for 30 seconds, then bonded at 75°C for 1 hour. Assembled devices were plasma treated as described for 1 minute and a solution of 10 μg mL^-1^ bovine plasma fibronectin (Sigma) in water was directly added to seeding ports. After 1 hour, devices were washed with PBS and stored overnight at 4°C.

### Cell treatments

Vinblastine sulfate (Sigma-Aldrich Corp., St. Louis, MO) and paclitaxel (Sigma) were stored in stock concentrations of 1 mM in DMSO, Y-27632 dihydrochloride (Sigma) was stored as 15 mM stock in water, cyclo-RGD(fV) peptide (Enzo Life Sciences Inc., Farmingdale, NY) was stored at 4 mM in PBS, and latrunculin A (Sigma) was stored at 0.5 mM in DMSO. All drug stocks were kept frozen at −20°C. DMSO volume added to dishes did not exceed 0.1% in any condition.

### Cell culture and seeding in microchannel devices

Human U251 glioma cells were cultured in DMEM/F-12 media (Gibco, ThermoFisher Scientific, Waltham, MA) containing 8% fetal bovine serum (Gibco) and 1% penicillin/streptomycin (Corning, Inc., Corning, NY). Cells were cultured in T25 flasks in an incubator maintained at 37°C and 5% CO_2_ and passaged using 0.25% trypsin-EDTA (Corning). Time-lapse movies were acquired in complete media without phenol red. U251 cell line origin was verified in a previous publication (25). For expression of fluorescent fusion proteins, cells were transfected using FuGENE^®^ HD transfection reagent (Promega Corporation, Madison, WI) according to manufacturer instructions at least 24 hours prior to seeding cells in devices. Fluorescent proteins expressed in this way included eGFP-β-actin (gift from Paul Letourneau, University of Minnesota) and EB1-eGFP (gift from Lynne Cassimeris, Lehigh University; (43)).

Devices were pre-incubated in media (containing drugs or DMSO, if appropriate) at 37°C for at least 1 hour prior to cell seeding. Cells were detached from flasks using 0.25% trypsin-EDTA (Corning), re-suspended in culture media, and nuclei were labeled using a nucleus counterstain (NucBlue Live, ThermoFisher) according to manufacturer instructions. Following labeling, cells were re-suspended in media to a density of 20×10^6^ cells per mL. Media was removed from device dishes and seeding ports, and a 5 μL aliquot of cell suspension (10^5^ cells) was added to each seeding port. Devices containing cells were returned to the incubator for 30 minutes before adding sufficient media to cover the device inlets.

### Time-lapse imaging of cells in microchannels

Time-lapse images were acquired on either a Nikon TiE or Nikon Ti2 epifluorescence microscope under control of NIS Elements software (Nikon Instruments Inc., Melville, NY). Each system is outfitted with a humidified environment containing 5% CO_2_ and stable temperature control at 37°C, provided by a stagetop incubator (OkoLab Llc., Ottaviano, Italy), a ProScan III motorized stage (Prior Scientific Inc., Rockland, MA), a white light transmitted LED (CoolLED Ltd., Andover, UK), LED fluorescence illumination (Spectra X; Lumencor, Beaverton, OR), and DAPI/FITC/TxRed filter set (Chroma, Cat#89014). The TiE system is equipped with a Zyla 4.2 sCMOS camera (Andor Technologies Ltd., Belfast, UK) and the Ti2 stand is equipped with a Zyla 5.5 sCMOS camera (Andor). Time-lapse images of device regions containing migrating cells were acquired in the transmitted, DAPI, and FITC (if expressing GFP-tagged proteins) channels every 5 minutes over 8-18 hours using a 20x/0.45NA phase I lens with 2×2 pixel binning (645 nm spatial sampling).

### Confocal z-stacks of cells in channels

Confocal z-stacks were acquired on an LSM7 LIVE swept-field confocal microscope (Carl Zeiss Microscopy, LLC; Jena, Germany) using a 40x/0.95 NA plan-apofluor objective and Optovar set to 1x magnification. The 488 nm and 405 nm lasers and BP 520-555 and BP 415-480 filters enabled simultaneous acquisition of the blue and green channels. Images were digitized using a 512 pixel 11 bit linear detector. Zen imaging software (Zeiss) provided system control during imaging.

### Semi-automated tracking of individual cell nuclei

Nucleus position and shape were obtained from the DAPI channel of image sequences using custom semi-automated image segmentation and tracking Matlab scripts (MathWorks) as previously described for tracking fluorescently labeled cells migrating in *ex vivo* brain slice cultures (24). Image sequences were cropped to include a single cell that was not in contact with another cell, as ascertained from phase contrast images. Skipped frames (e.g. due to contact with another cell or object in the channel) were omitted during analysis. Centroid coordinates were used to calculate MSD using the overlap method (40) and analyzed in the same way as 1D CMS data. Individual cells in some conditions were analyzed using a PRW model (13). PRW model fits were performed in the same way as fits to the diffusion-convection model.

### Statistical analysis

Experimental data represent accumulated measurements from N ≥ 2 biological replicates for each condition. Numbers of individual simulations are given in the figure legends. All statistical analysis of experimental or simulated data was performed using the Matlab Statistics Toolbox (MathWorks). We used the non-parametric Kruskal-Wallis test, since the data did not typically follow a normal distribution. When a comparison included more than two groups, we corrected multiple pairwise comparisons using a Dunn-Sidăk *post hoc* test. Statistical significance for a particular comparison was reached when p < 0.01.

## Results

### Cell migration simulator predicts persistent or random migration as a function of cell polarity

In the previously described CMS (24, 25), each protrusion module functions as a copy of the motor-clutch model (23), with myosin II motor proteins generating F-actin retrograde flow, while the transient binding of molecular clutches establish traction forces required for motion. Protrusion modules elongate by actin polymerization, while protrusion nucleation and capping – a process that stochastically prohibits actin polymerization – govern protrusion turnover and cell polarization. Mass conservation of cellular components (e.g. motors, clutches, actin) enforces their distribution between modules and a maximum cell length. With only modest complexity (18 parameters to describe the cell plus a compliant substrate spring with variable stiffness, κ_sub_; all parameters summarized in **Table S1**), the CMS predicts glioma cell migration in environments of varying stiffness and adhesiveness (24, 25). However, as these studies used parameter sets calibrated for cells on 2D substrates, it is unclear how well CMS predictions hold for cells migrating in confined channels.

We first sought to determine which parameter changes or modifications to the CMS were necessary to recapitulate persistent migration that is often experimentally observed in channels (9–11). To simulate migration in a confined 1D channel that limits protrusions perpendicular to the channel axis, we modified the CMS to solve for cell coordinates only within a single spatial dimension (**Figure 1A**). We introduced cell polarity as an additional variable ψ, which represents the probability that a new module will be generated in the +x direction (as described in **Materials and Methods**). Increasing ψ led to increasingly persistent motion in the +x direction (**Figure 1B**), corresponding with non-linear mean-squared displacement (MSD) as a function of time lag (**Figure 1C** and **D**). Simulations without a bias in module nucleation direction (ψ = 0.5) had approximately linear MSD versus time curves (**Figure 1C**), reminiscent of random walk behaviour (40). Intriguingly, changing ψ did not change the average number of protrusions (**Figure S1** in the **Supporting Material**), confirming that persistent migration arises due to directionally biased nucleation and turnover. Through changing polarization state, 1D CMS can thus replicate persistent or random motility, providing an opportunity to test model predictions against data obtained in 1D experimental conditions.

### Human glioma cells move persistently in confined microchannels

To develop an assay for glioma cell migration that includes both 1D spatial guidance cues and mechanical confinement, we used photolithography and PDMS molding techniques to manufacture microchannel devices. Devices were designed with 12 μm wide channels with 5 μm height (60 μm^2^ cross sectional area) around a central seeding port (**Figure 2A** and **B**). We used PDMS replica-molding techniques to create microfluidic devices containing the channels, bonded the devices to glass bottom dishes (**Figure 2A**) and functionalized channels with fibronectin (41).

**Figure 2.**
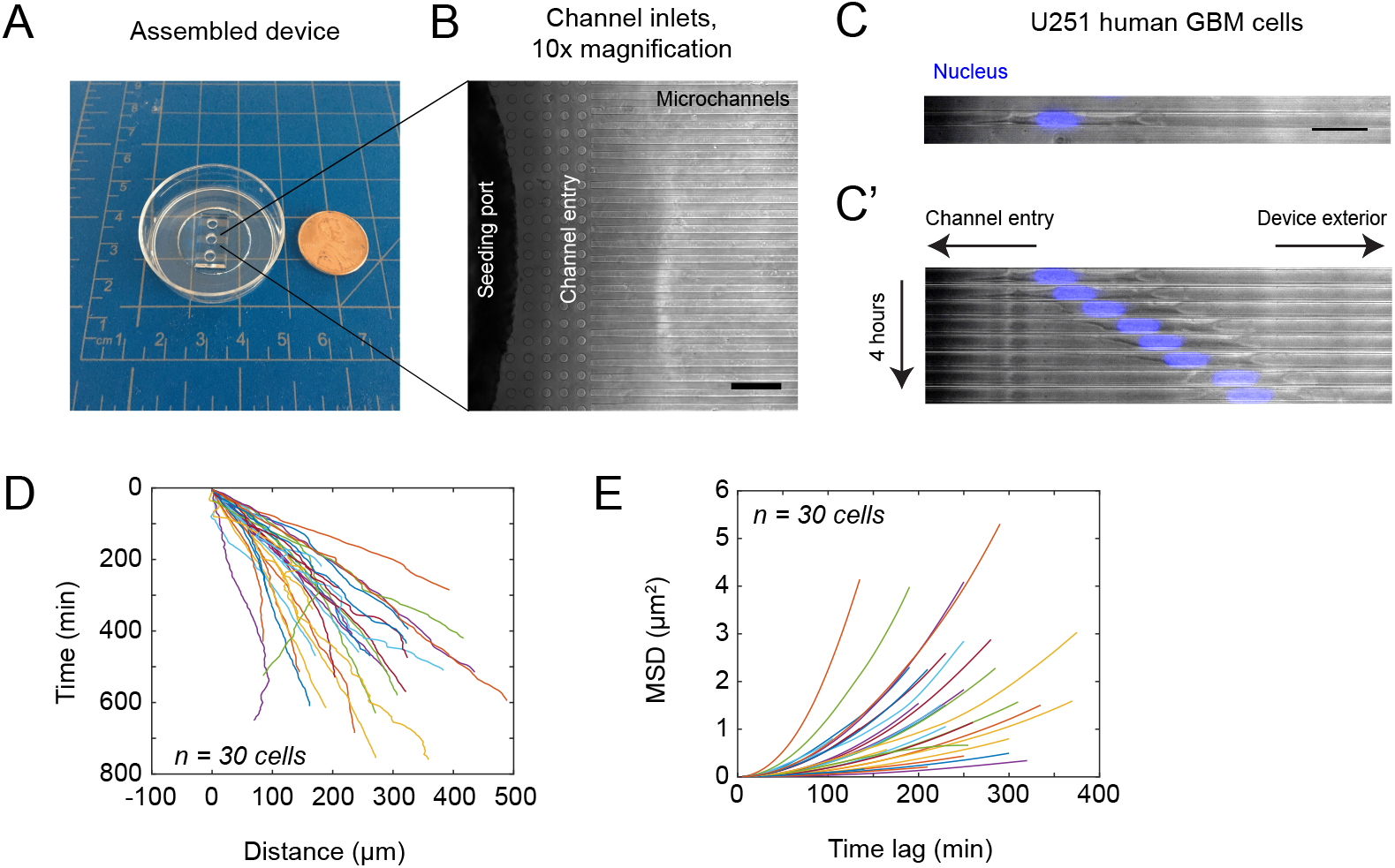
Single cell nucleus tracking method to analyze glioma cell migration in confined microchannels. (A) Photograph of an assembled device bonded to a 35 mm glass bottom dish. Note channels extending from drilled seeding ports to the exterior of the device. A US one-cent coin is shown for scale and the grid spacing is 1 cm. (B) Image of drilled seeding port showing the entry chamber and 12 μm wide channels. Image was acquired at 10x magnification using phase contrast optics. Scale, 100 μm. (C) U251 glioma cell migrating in a confined 12 μm-wide channel imaged using phase contrast and fluorescence. Nucleus counterstain is shown in blue. Images acquired at 20x magnification. Scale bar 50 μm. (C’) Time-lapse sequence of images acquired for the cell in panel B. Images are displayed at 30-minute intervals. (D) Nucleus x-position versus time as measured for n = 30 individual cells from a single control experiment. Coordinates are plotted relative to the initial tracking position for each cell, such that x(τ = 0) = 0. For display purposes, coordinates of cells moving in the –x direction were reversed. (E) MSD versus time lag for the individual cells in panel E. For clarity, error bars are not shown for individual traces.

We generated time-lapse movies of cells moving in channels (**Figure 2C** and **D**) and tracked fluorescently labeled nuclei using a previously described semi-automated tracking method (24). U251 cells spontaneously entered the channels from seeding ports, even in the absence of an established chemokine gradient, as is often used in microfluidic migration assays (11, 28). Cells elongated along the channel axis, typically forming long leading protrusions that tracked along either or both channel walls, while trailing protrusions were typically smaller (**Figure 2D** and **Movie S1** in the **Supporting Material**). The nucleus typically filled the lateral width of the channel (**Figure S2** in the **Supporting Material**), and maintained a near-constant size and shape. Cells moved persistently along the channel axis towards the device exterior.

We calculated MSD versus time lag for experiments in a similar way as with simulations. MSD versus time plots (**Figure 2E** and **F**) were nonlinear with concave up character, similar to the 1D CMS with high polarity (e.g. ψ = 0.9). However, while simulation tracks contain frequent, short-term direction switches within each individual cell position (**Figure 1B**), cell position traces from experiments were smoother (**Figure 2E**). One possible explanation for this difference is that, in 1D CMS, the stiff substrate causes some slippage of the cell body clutches (since the simulated cells are well above their optimal stiffness (25)), leading to the short timescale reversals seen in cell position. A few cells exhibited “start and stop” motion as observed in GBM cells migrating in brain slice cultures (6, 24) and complete directional reversals were rare.

### A diffusion-with-drift model, rather than a persistent random walk model, describes cell migration in microchannels

The random walk model that we have used in previous analysis of 2D migration (25) would produce poor fits to our experimental data because persistent motion results in a nonlinear relationship between MSD and time lag. We have previously used a persistent random walk (PRW) model to describe U251 cell migration on single nanometer-scale suspended fibers (13). In the PRW model (Eqn. 9) the quantity 〈*r*^2^(*t*)〉 represents MSD, n is the spatial dimension (n = 1 for a 1D fit), and t represents time lag. Cell speed (S) and persistence time (P) are the two fitting parameters.

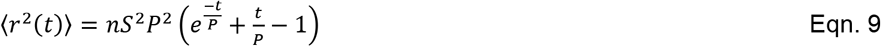

PRW model fits to individual control cell trajectories (n = 403 cells from 12 experiments) yielded a mean speed of S = 0.74 ± 0.05 μm min^-1^ (**Figure S3** in the **Supporting Material**). Comparing to other studies in confined channels of similar dimension, mean speed was comparable to other tumor cell lines (8, 10, 11) and mesenchymal stem cells (19), but much slower than immune cells (9, 18, 41). However, persistence times obtained for individual cells were up to two orders of magnitude longer than the maximum experimental duration of 18 hours, or 10^3^ minutes. In order to avoid having to artificially constrain fits, we sought an alternative model to describe the data.

A 1D diffusion-convection model is based off of the random walk model for migration (40) which describes a linear relationship between MSD and time lag that are used to calculate a motility coefficient, μ. To account for persistent motion that results in a nonlinear relationship between MSD and time lag, a quadratic, time-dependent term represents the velocity, v.

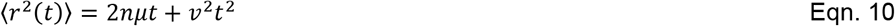

Eqn. 10 produced a reasonable fit to the pooled MSD from all control data (n = 403 cells, N = 12 replicates; **Figure 3A**). Individual cell trajectories revealed large variability in both motility (**Figure 3B**) and velocity (**Figure 3C**). We also analyzed MSD curves obtained from the 1D CMS data obtained with varying ψ (**Figure 3D**). Very little difference was observed in motility between the three values of ψ (**Figure 3E**); however, as expected, v was smallest for the ψ = 0.5 data (**Figure 3B**). This confirms that Eqn. 10 can capture random motility behavior for small values of v, whereas otherwise the quadratic term dominates at long timescales. Interestingly, the overall mean motility coefficient was larger for simulations (**Figure 3E**) than experiments (**Figure 3B**). Simulated cell velocities (**Figure 3F**) were similar to experiments (**Figure 3C**), and in turn all were in close agreement with values obtained using the PRW model (v_exp_ = 0.51 ± 0.02 μm min^-1^ or v_exp_ = 8.5 ± 0.3 nm s^-1^). We conclude that the 1D diffusion-convection model can capture a wide range of migration behaviors for cells migrating on aligned structures. We also note that our results suggest that caution should be used in applying the PRW model to interpret data collected for cells on 1D aligned substrates, as it can produce aberrant parameter values that lie outside the bounds of the experimental system.

**Figure 3.**
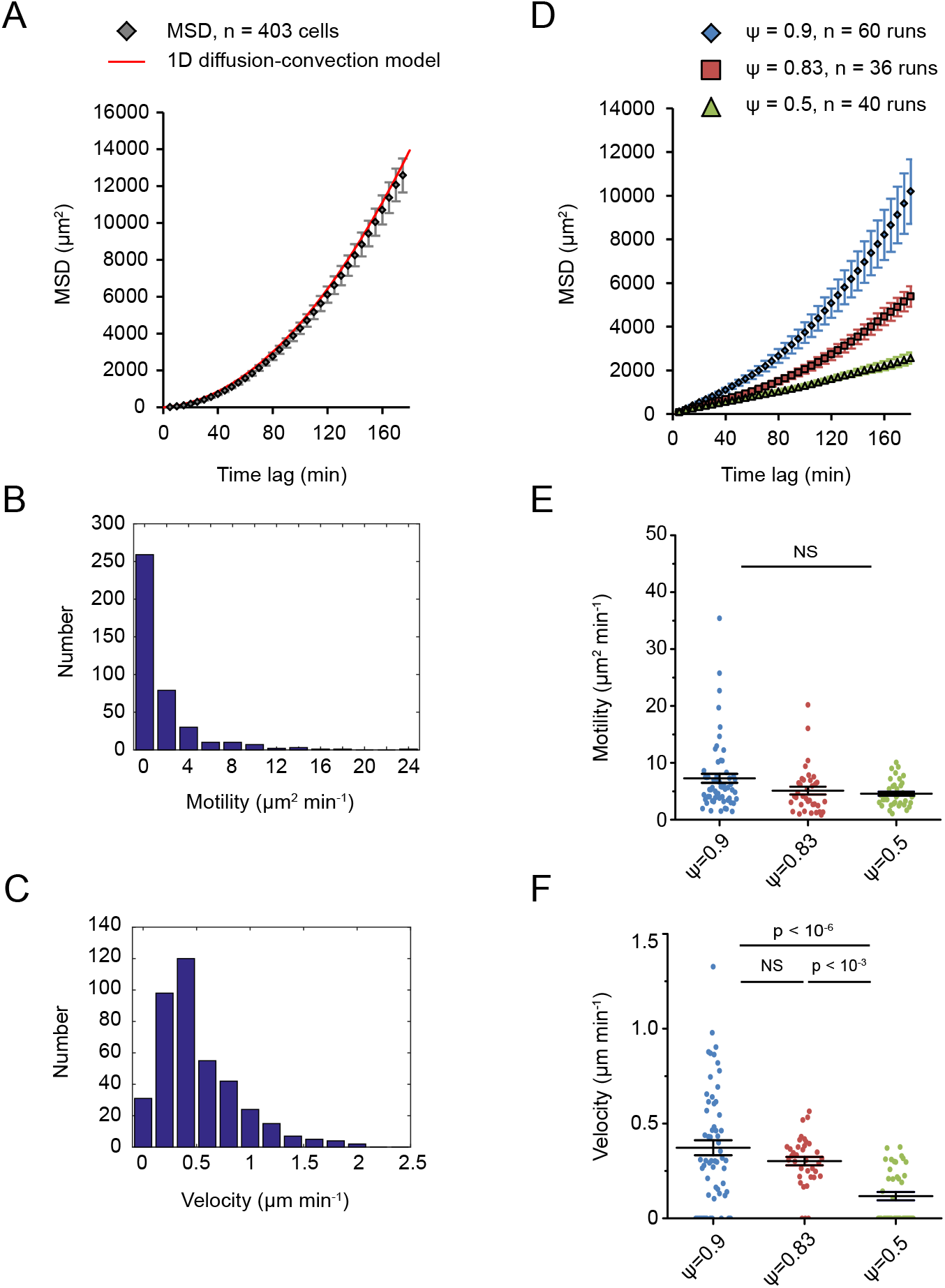
Analysis of experimental and simulated confined cell trajectories using a diffusion-convection model. (A) MSD versus time lag for all control experiments: n = 403 cells pooled from 12 independent experiments. Data points are mean ± SEM, red line is a least squares minimization fit to a 1D diffusion-convection model. Fit parameters: μ = 2.83 μm^2^ min^-1^ and v = 0.62 μm min^-1^. (B) Histogram of motility coefficients from individual cell trajectories: μ = 1.61 ± 0.14 μm^2^ min^-1^ (mean ± SEM). (C) Histogram of velocities for the cell trajectories in panel B: v = 0.51 ± 0.02 μm min^-1^ (mean ± SEM). (D) Mean of MSD versus time lag for the 1D CMS with ψ = 0.9, 0.83, or 0.5. Data are presented as mean ± SEM. (E) Motility coefficients for the individual simulation trajectories shown in **Figure 1**. (F) Velocities for the individual simulations in panel C. All motility coefficients and velocities are measured from individual fits to a diffusion-convection model (Eqn. 10). Pairwise p-values obtained by Kruskal-Wallis test with Dunn-Sidăk *post hoc* test. NS indicates no significant difference between groups, defined as p > 0.01.

### Microchannel based migration requires integrin-mediated adhesion and myosin II force generation

U251 cell migration on 2D hydrogel substrates involves both myosin II activity and integrin-mediated adhesion (25). Since the 1D CMS can predict migration trends in control cells, we sought to test the robustness of model predictions in microchannels by introducing pharmacological inhibitors of cell components involved in 2D migration. Interestingly, changing clutch number (n_clutch_) in simulations produced a biphasic trend in motility and velocity (**Figure 4A** and **B**), where both quantities are largest for an intermediate n_clutch_. These findings are consistent with earlier theoretical and experimental results (24, 44). We sought an experimental test of this finding in microchannels, so we treated cells with cyclo-RGD(fV) peptide (cRGD), which competitively inhibits α_v_β_3_ integrin binding to fibronectin (45). On 2D hydrogels, cRGD treatments produce effects consistent with decreasing clutch number (e.g. decreased traction force, increased F-actin flow, decreased cell spreading and migration; (25)). Treatment with 0.1-1 μM cRGD revealed biphasic trends in motility (**Figure 4C**) and velocity (**Figure 4D**) where intermediate doses modestly increased motility and velocity. Thus, U251 cell migration in channels is integrin-dependent and follows a well-known adhesiveness relationship predicted by biophysical theory (44).

**Figure 4.**
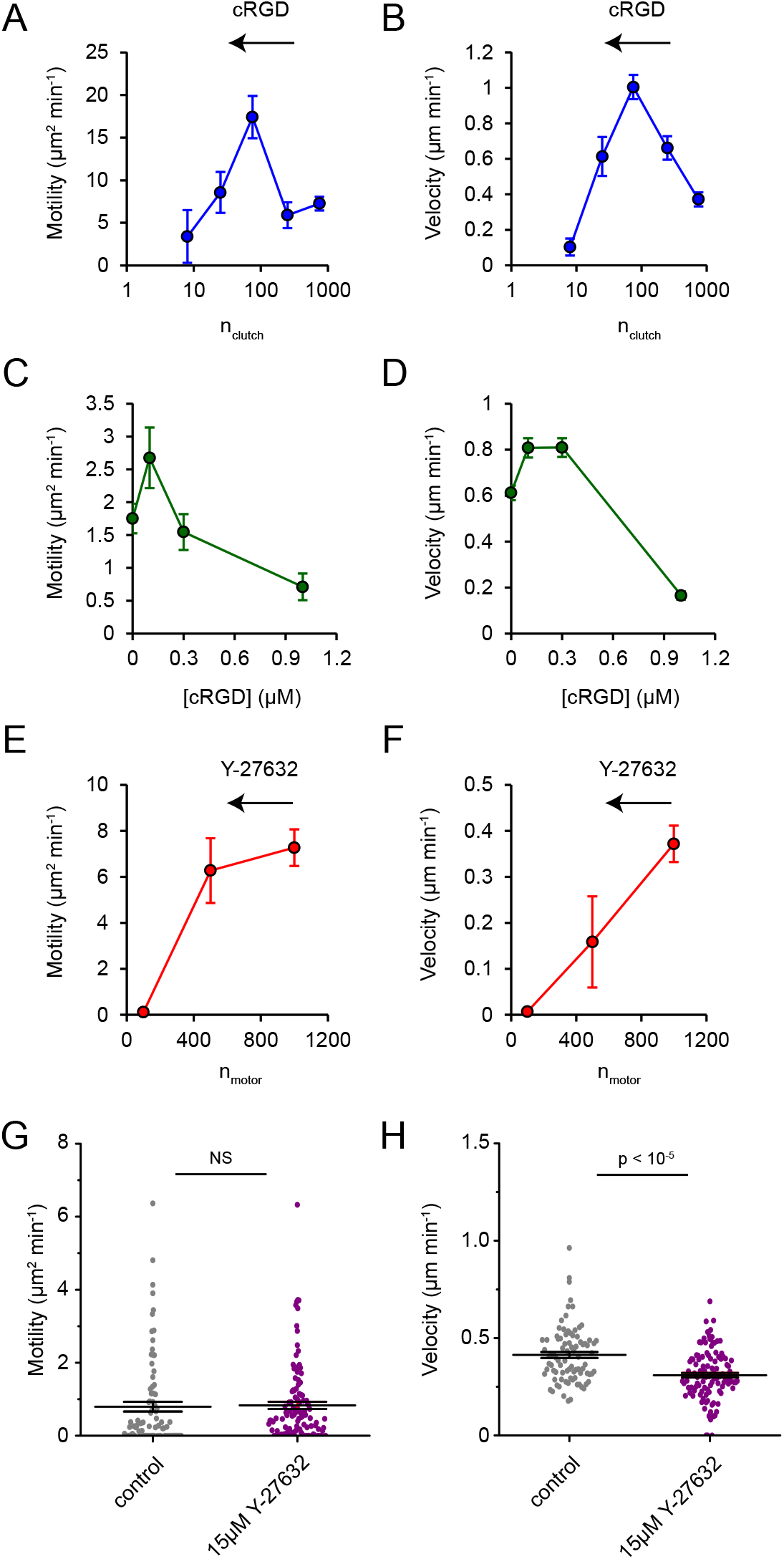
Confined glioma cell migration sensitivity to inhibitors of integrin-mediated adhesion and the ROCK pathway. (A) 1D CMS prediction of motility coefficient where n_clutch_ was varied independent of other parameters: n_clutch_ = 8, 25, 75, 250, 750, obtained with: n = 8, 8, 16, 16, 60 runs, respectively. All simulations had n_motor_ = 1000, other parameters are reported in **Table S1**. Statistics for pairwise comparisons are reported in **Table S2**. (B) Velocities for the fits in panel A. (C) Motility coefficients for U251 control cells, or cells treated with 0.1, 0.3, or 1 μM cRGD, n ≥ 72 cells per condition. (D) Velocities for the fits in panel B. (E) 1D CMS prediction of motility where n_motor_ was varied independent of other parameters: n_motor_ = 100, 500, 1000, collected from n = 60 runs for base parameters, n = 8 for all other conditions. All simulations had n_clutch_ = 750, other parameters are reported in **Table S1**. Statistics for pairwise comparisons are reported in **Table S2**. (F) Velocities for the fits in panel E. (G) Motility coefficients for U251 control cells, or cells treated with 15 μM Y-27632, n = 87, 112 cells. (H) Velocities for the fits in panel G. Error bars in all panels represent mean ± SEM. NS indicates no significant difference between groups, defined as p > 0.01. All motility coefficients and velocities are measured from individual fits to a diffusion-convection model (Eqn. 10).

Myosin II inhibition slows glioma cell migration on 2D hydrogels (25) and *ex vivo* brain slices (6, 46). Reducing myosin II motor number (n_motor_) as an independent parameter nearly monotonically reduced motility (**Figure 4E**) and velocity (**Figure 4F**). Direct inhibition of myosin II with blebbistatin is not compatible with our nucleus tracking method, since imaging at near-ultraviolet wavelengths is known to inactivate the drug and cause conversion to cytotoxic species (47). Y-27632 is an inhibitor of Rho-associated kinase (ROCK), which is one of the major pathways for myosin II activation (48), so we used this as an orthogonal approach to reduce myosin II activity. In U251 cells, Y-27632 (15 μM) had little effect on motility (**Figure 4G**), but reduced velocity (**Figure 4H**). These results are consistent with 1D CMS predictions of minimal reduction in n_motor_ (**Figure 4E** and **F**). Weak sensitivity to ROCK inhibition in confinement in other cell types (11, 19, 49) suggests other pathways to activate myosin II or alternative modes of force generation.

### Glioma cells require actin polymerization dynamics during microchannel-based migration

Several studies suggest that the roles of the actin and microtubule cytoskeleton in migration change in confined environments, compared to cells on flat 2D substrates (11, 19, 29). For example, breast carcinoma and sarcoma cells migrating in confinement via an osmotic engine mechanism are insensitive to inhibitors of actin polymerization (29). We tested this prediction in our system by treating U251 cells with latrunculin A (LatA), which is a potent actin polymerization inhibitor in sub-micromolar doses (50). Compared to vehicle controls (DMSO), 50 nM LatA slowed migration, while addition of 500 nM LatA nearly completely stalled motion (**Figure 5A** and **B** and **Movie S2** in the **Supporting Material**). Simulating the effects of intermediate LatA doses by reducing v_actin,max_ (the maximum actin polymerization rate) reduced cell motility and velocity (**Figure 5C** and **D**). Interestingly, the effect of decreasing v_actin,max_ was much milder than our earlier reports in 2D (26). Alternatively, because LatA prevents actin subunits from binding to F-actin barbed ends (50), we tested the possibility that increased module capping could recapitulate our results. Capping occurs at a rate k_cap_, and stochastically terminates actin polymerization to facilitate module turnover. Increasing k_cap_ 10-fold reduced simulated motility and velocity (**Figure 5C** and **D**), consistent with our predictions. These accumulated results suggest that actin protrusion and turnover rates regulate migration speed in the 1D CMS, and argue against U251 cells employing osmotic pressure-based migration in our devices.

**Figure 5.**
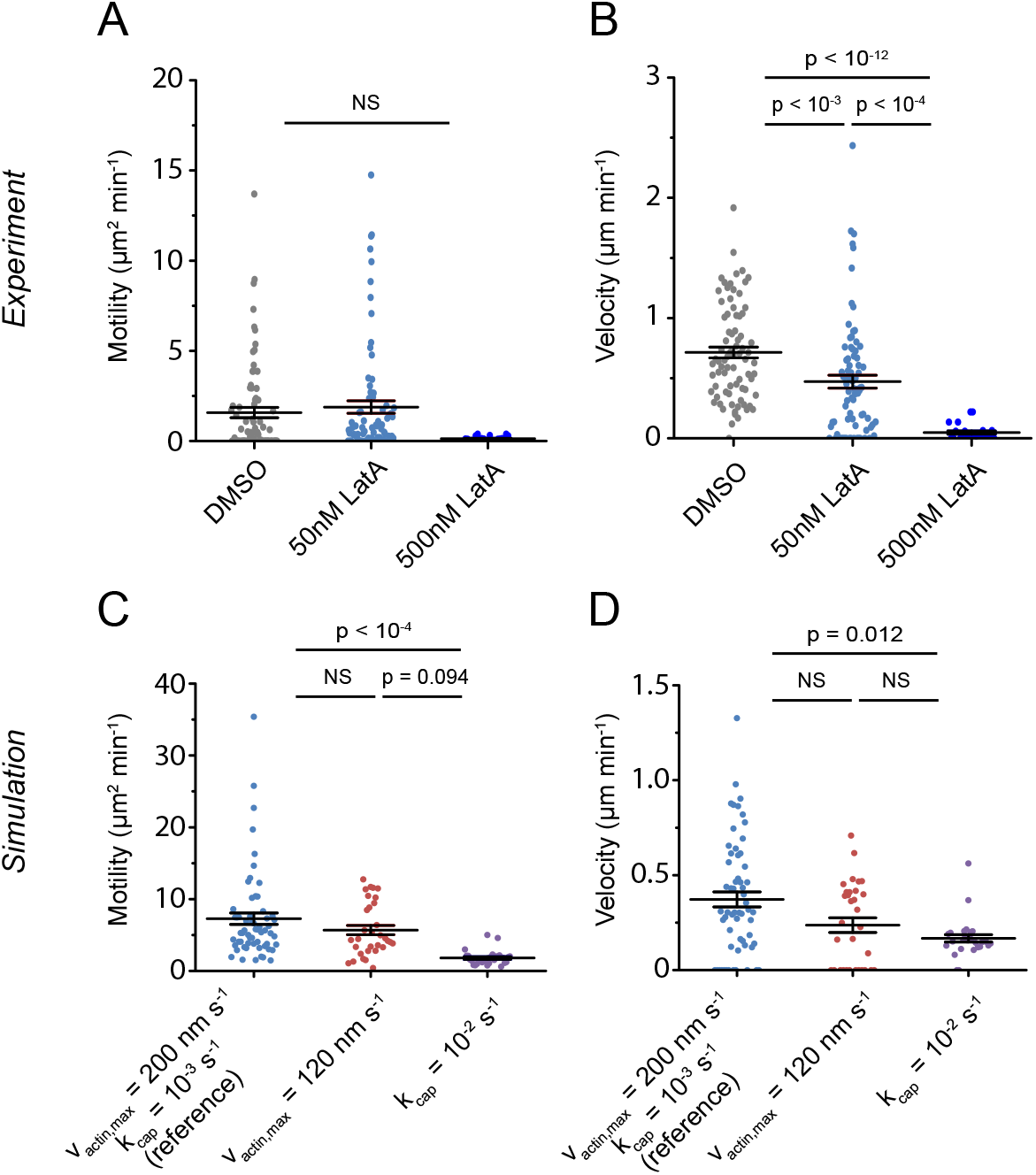
Confined glioma cell migration sensitivity to inhibitors of actin polymerization dynamics. (A) Motility coefficients for U251 cells transfected with eGFP-β-actin and treated with vehicle control (DMSO), 50 nM, or 500 nM LatA, n = 78, 81, 23 cells. (B) Velocities for the cells in panel A. (C) Motility coefficients for 1D CMS with a reference parameter set (v_actin,max_ = 200 nm s^-1^, k_cap_ = 0.01 s^-1^, all other parameters reported in **Table S1**), or with v_actin,max_ = 120 nm s^-1^ or k_cap_ = 0.01 s^-1^ altered as single parameter changes, n = 60, 28, 24 runs. (D) Velocities from the fits in panel C. Simulations with base parameters are reproduced from **Figure 1**. Error bars are mean ± SEM. All motility coefficients and velocities are measured from individual fits to a diffusion-convection model (Eqn. 10). Pairwise p-values obtained by Kruskal-Wallis test with Dunn-Sidăk *post hoc* test. NS indicates no significant differences between groups, defined as p > 0.01.

### Microtubule-targeting agents slow glioma cell migration consistent with increasing module nucleation in simulations

Dynamic microtubules play key roles in establishing and maintaining the polarity of migrating cells (34). Two main classes of MTAs promote either microtubule polymer assembly or disassembly; yet the common effect of each is kinetic stabilization or attenuation of self-assembly dynamics (26, 51). Treating U251 cells with either PTX or VBL (each at 100 nM) significantly slowed migration velocity within the channels, compared to vehicle control (**Figure 6A** and **B**). One of the most pronounced effects of kinetic stabilization is the loss of end-binding protein 1 (EB1) accumulation on microtubule plus-ends (26, 51). Balzer *et al*. reported pronounced effects of MTAs on cells in confinement, and reported that arrival of EB1-decorated microtubule ends was concomitant with forward protrusion of the leading edge (11). This correlation suggests that kinetic stabilization reduces the rate of microtubule delivery to the leading edge, which in turn slows progressive forward migration along the channel.

**Figure 6.**
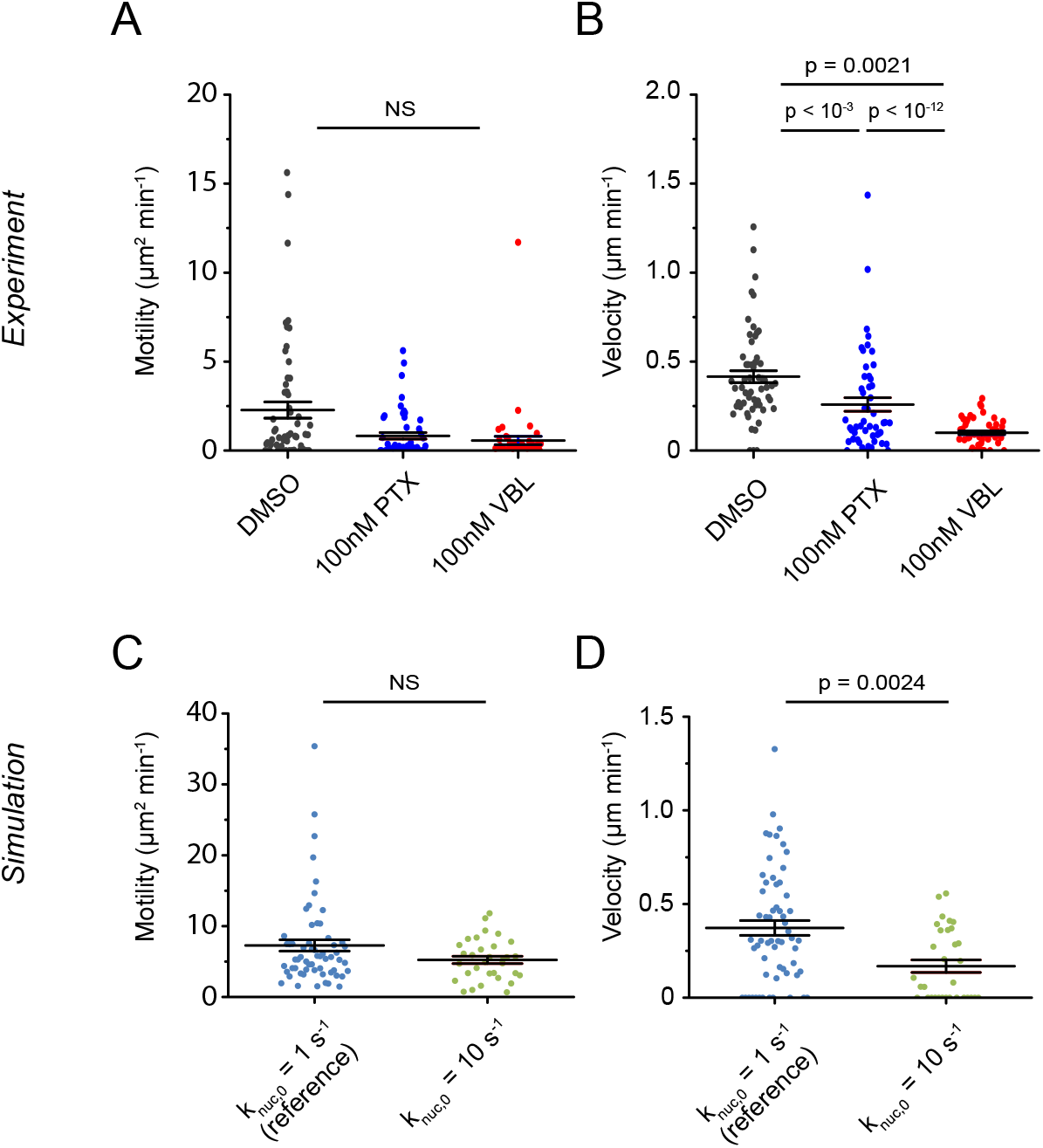
Microtubule-targeting agent effects on confined glioma cell migration and relation to module nucleation rate. (A) Motility coefficients for U251 cells treated with vehicle control (DMSO), 100 nM PTX, or 100 nM VBL, n = 58, 52, 50 cells. (B) Velocities from the fits in panel A. (C) Motility coefficients for 1D CMS with a reference parameter set (k_nuc,0_ = 1 s^-1^, all other parameters reported in **Table S1**), or k_nuc,0_ = 10 s^-1^ altered as a single parameter, n = 60, 28 runs. (D) Velocities from the fits in panel C. Simulations with base parameters are reproduced from **Figure 1**. Error bars are mean ± SEM. All motility coefficients and velocities are measured from individual fits to a diffusion-convection model (Eqn. 10). Pairwise p-values obtained by Kruskal-Wallis test with Dunn-Sidăk *post hoc* test. NS indicates no significant differences between groups, defined as p > 0.01.

Our earlier study (26) concluded that kinetic stabilization slows U251 cell migration on 2D substrates by either slowing actin polymerization (as in **Figure 5C** and **D**) or increasing the basal nucleation rate of new protrusions (k_nuc,0_) through microtubule-dependent signaling mechanisms. Each of these changes caused an increase in the number of protrusion modules in 2D, which correspondingly slowed migration. However, it is unclear how these predictions would hold for cells in a confined 1D channel. Increasing k_nuc,0_ reduced simulated cell migration (**Figure 6C** and **4D**), suggesting that MTAs inhibit glioma cell migration by similar mechanisms in 1D confinement as they do in 2D (26). Moreover, we find that parameter values and conclusions drawn from 2D studies can be rigorously tested for mechanistic relevance in other environments. Altogether, our results suggest the 1D CMS can be used to accurately test biophysical mechanisms of tumor cell migration as well as the effects of novel or existing anti-motility chemotherapies.

## Discussion

Glioma cells migrate in conditions of mechanical confinement within their major invasion routes in the brain (5, 6). Micron-scale PDMS channels can reproduce the mechanical confinement found in aligned brain tissue structures in a highly reproducible fashion. We find that CMS predictions are consistent with spontaneous cell migration behaviors in channels with minimal changes from parameters previously calibrated on 2D hydrogel substrates (25). Furthermore, parameter changes derived from earlier studies on 2D substrates (26) can reproduce the effects of various motility-altering drugs, suggesting that glioma cells use the same motor-clutch machinery in these two different environments.

Recent physics-based models predict confined migration phenotypes that do not fit the classical description of cell migration. One particularly interesting observation is the variable influence of substrate adhesiveness, whereby cells can move by non-specific friction between cortex and channel walls, rather than relying on integrin-mediated adhesion (21, 27). The 1D CMS similarly predicts a regime where weakening adhesion by reducing n_clutch_ could actually increase cell migration velocity (**Figure 3A** and **B**), which we observed experimentally (**Figure 3C** and **D** in this study and in (24)). These results are consistent with a biphasic adhesiveness relationship predicted by previous biophysical theory (44) and the current study, where migration is fastest at an intermediate adhesiveness. Liu *et al*. (27) found that fast, polarized migration occurred when cells were confined between surfaces functionalized to promote non-specific adhesive interactions (e.g. frictional forces), while most cells moved very slowly on unconfined substrates treated the same way. Friction-based adhesive forces could be specifically incorporated in the motor-clutch framework as clutches with high binding and unbinding rates (k_on_ and k_off_, respectively) and low bond forces (F_bond_). Increasing the absolute number of clutches (n_clutch_) to represent the increase in adhesiveness provided by the second surface in contact with the cell, could shift cells to the intermediate “fast motility” regime, consistent with the results of Liu *et al*.

Stroka *et al*. (29) describe tumor cell movement in confined channels driven by intracellular osmotic pressure. In this osmotic engine model, asymmetrically distributed ion pumps at the leading and trailing edge generate osmotic pressure to create a net force on the semi-permeable plasma membrane that propels the cell forward. Extensive contact between the cortex and the channel wall provides the means of force transmission. We find that U251 cell migration in microchannels is sensitive to LatA (**Figure 4A** and **B**), contrasting the experimental observation that osmotic pressure-based migration does not require actin polymerization (29). There are two non-exclusive explanations for these contrasting findings. First, the channels in the present study have a larger cross sectional area than the ones used by Stroka *et al*. (30 μm^2^ in (29) compared to 60 μm^2^ in this study). Narrow channels increase the hydraulic resistance that cells must overcome, which could compete with adhesion-dependent migration (30). Second, osmotic effects could be cell type-specific as not all cells display actin-independent motility in a high degree of confinement (19). This is possibly related to the expression of particular ion channels (29). Regardless, osmotic pressures could be included as a modification to the 1D CMS by including an additional outward force that facilitates actin polymerization at the module end. These modifications would possibly require the 1D CMS to include an explicit membrane tension term, which could also be pursued in future studies.

### Advantages and limitations of 1D CMS

A limitation of the current 1D CMS is that it relies upon an empirical polarity parameter (ψ) to replicate experimental results, rather than enabling symmetry breaking by random module nucleation (24, 25). Previous studies have employed similar mathematical expressions for cell polarity. DiMilla *et al*. coupled an intracellular transport model to their mechanical model, which caused simulated cells to accumulate adhesion receptors at the cell front (44). Pathak and Kumar’s model relied upon a traction polarization term that increased in narrow channels, based on stress fibers and active myosin II aligned along the channel axis (52). In the present study, we reasoned that U251 cells polarize upon channel entry from the seeding ports and maintain a near constant polarity during migration, since directional reversals were rare (**Figure 2A** and **B**). 1D CMS predicts that parameters regulating actin dynamics (e.g. capping, nucleation, and polymerization) may antagonize cell polarity, possibly through cross talk with dynamic microtubules. In 1D CMS, either reducing v_actin,max_ or increasing k_cap_ could be responsible for this effect, as either parameter change slows simulated cells (**Figure 5C** and **D**).

### Advantages and limitations of PDMS microchannels

Photolithography and PDMS replica molding of channel structures provides excellent, reproducible spatial control over channel dimension, allowing medium-high throughput individual analysis of migration phenotypes (~1,000 cells in a single study). Controlled and reproducible feature size is a distinct advantage over 3D collagen gels, where it is difficult to independently control pore size and stiffness. Although the current study uses devices with stiff PDMS walls (Young’s modulus ~ 1000 kPa (53)) and glass dishes, mixing two-component PDMS elastomer in different ratios can yield substrates with significantly lower Young’s moduli (53). Some studies bridge two disparate stiffness regimes by independently measuring cellular properties in both collagen gels and stiff microfluidic devices (10). A complication with this approach is that a number of other parameters besides stiffness and pore size could change between the two assay types (e.g. fiber alignment, remodeling, adhesive contact area), each of which could independently influence migration.

Pathak and Kumar used similar photolithography techniques to create tracks of varying width (10-40μm) in compliant polyacrylamide hydrogels (Young’s modulus = 0.4-120 kPa) enabling them to independently control lateral confinement and mechanics (52). Interestingly, they reported biphasic glioma migration speeds as a function of stiffness, which is a key prediction of the CMS in 2D (25). Biphasic speed as a function of adhesiveness *in vitro* (this study) and in tissues (24) also support our conclusion that glioma cells use a motor-clutch mechanism in confined environments. However, while this approach creates a versatile platform with tunable modulus and dimension, cells are not enclosed in channels as in our devices, or in tissue. It is also unclear whether polyacrylamide molding techniques could generate ~1-2 μm width constrictions, which are necessary to impair nuclear passage for many cells (10, 18, 28) and can be reproducibly created in PDMS devices (18, 41).

### Connection to GBM tumor progression and therapy

Using cell parameters calibrated from previous 2D studies (25), we obtained reasonable theoretical predictions of experimentally measured migration speeds of ~8 nm s^-1^ or 25 cm yr^-1^, within the range of GBM tumor growth rates measured in the clinic (54). Since osmotic migration effects have been suggested as a mechanism for GBM progression (4), it will be interesting to further test whether GBM cells migrating *in vivo* primarily use actin-based (24) or osmotic pressure-based migration (29), since these are predicted to occur in different regimes of confinement (e.g. varying channel geometries (30)) that could correspond to different brain regions. Measuring cell migration and proliferation rates in tissue-relevant environments can provide input data for mathematical models of tumor progression and treatment (54–56) and could be used to predict how cellular responses to chemotherapy influence tumor dispersion. The combined simulation and experimental framework presented in this paper represent a step towards predicting cell migration in confined tumor microenvironments and understanding mechanisms of GBM tumor invasion.

## Author Contributions

Conceptualization, L.S.P., M.P., and D.J.O.; Methodology, all authors; Software, L.S.P.; Validation, L.S.P. and M.R.S.; Formal Analysis, L.S.P., M.R.S., and D.J.O.; Resources, M.P. and D.J.O.; Writing – Original Draft, L.S.P. and D.J.O.; Writing – Review & Editing, all authors; Visualization, L.S.P.; Supervision, P.V., M.P., and D.J.O.; Funding Acquisition, L.S.P., M.P., and D.J.O.

## Acknowledgements

Thanks to members of the Odde and Piel laboratories for helpful discussions, and thanks to Prof. Margaret Titus for comments on the manuscript. The authors thank Dr. Rafaele Attia and Mark Fisher for their assistance with device design, as well as Geneva Doak, Jose Valdez, Dr. Daniel Lu, and Prof. David Wood for their assistance with photolithography and microfabrication. The authors thank Dr. Ana-Maria Lennon-Duménil for use of her laboratory space, and the Bioimaging Cell and Tissue Core Facility of the Institut Curie (PICT-IBiSA) for use of microscopes during pilot data collection. Simulations were run in part on high-performance computing resources at the Minnesota Supercomputing Institute. Portions of this work were conducted in the Minnesota Nano Center, which is supported by the National Science Foundation through the National Nano Coordinated Infrastructure Network (NNCI) under Award Number ECCS-1542202. Other funding support was provided by a 3M Science & Technology Fellowship (to L.S.P.), National Science Foundation Graduate Research Fellowship 0039202 and GROW program grant (to L.S.P.), a STEM Chateaubriand Fellowship (to L.S.P.), the University of Minnesota UROP (to M.R.S.), Association Nationale pour la Recherche grant ANR-16-CE13-0009 (to P.V.), European Research Council consolidator grant 311205 PROMICO (to M.P.), and National Institutes of Health grants U54 CA210190 and R01 CA172986 (to D.J.O). P.V. is an INSERM investigator.

