## Supplemental Material for "Glioma cell migration in confined microchannels via a motor-clutch mechanism"

### Supporting Tables

**Table S1.** 1D CMS base parameter set. Related to **Figure 1**.

| **Parameter** | **Description** | **Range** |
|---|---|---|
| *Motor parameters* | | |
| $n_{motor}$ | Number of myosin II motors | 1,000 |
| $F_{motor}$ | Myosin II stall force | 2 pN |
| $v_{motor}$ | Unloaded (maximum) myosin II velocity | 120 nm $s^{-1}$ |
| *Clutch parameters* | | |
| $n_{clutch}$ | Number of molecular clutches | 750 |
| $F_{bond}$ | Characteristic slip bond force for clutches | 2 pN |
| $\kappa_{clutch}$ | Clutch spring stiffness | 0.8 pN $nm^{-1}$ |
| $k_{on}$ | Pseudo-first order binding rate for clutches to F-actin | 1 $s^{-1}$ |
| $k_{off}$ | Basal first-order clutch unbinding rate | 0.1 $s^{-1}$ |
| *Substrate parameters* | | |
| $\kappa_{csubstrate}$ | Substrate spring stiffness | 1,000 pN $nm^{-1}$ |
| *Cell body and actin parameters* | | |
| $k_{cap}$ | Module capping rate | 0.001 $s^{-1}$ |
| $k_{nuc,0}$ | Maximum module nucleation rate | 1 $s^{-1}$ |
| $v_{actin,max}$ | Maximum actin polymerization velocity | 200 nm $s^{-1}$ |
| $A_{total}$ | Total actin pool available for protrusions | 100 µm |
| $\kappa_{cell}$ | Cell spring constant | $10^4$ pN $nm^{-1}$ |
| $L_{cell}$ | Initial module length | 5 µm |
| $L_{min}$ | Minimum module length | 100 nm |
| $n_{clutch,cell}$ | Number of cell body clutches | 10 |
| $\psi$ | Variable cell polarity factor | 0.9* |

*Cell polarity factor ψ was varied from 0.5-0.9 to simulate different polarization states of the cell. In simulations, this parameter is given as a variable input p, such that $\psi=p/(p+1)$ defines the threshold for nucleation in the +x direction.

**Table S2.** Pairwise p-values for the experimental and simulation conditions involving inhibitors of integrins and the ROCK pathway. Related to **Figure 4**.

**Figure 6A**. 1D CMS motility ($n_{motor}$ = 1000)

| $n_{clutch}$ | 25 | 75 | 250 | 750 |
|---|---|---|---|---|
| 8 | 0.95 | 0.02 | 1 | 0.06 |
| 25 | | 0.01 | 0.98 | 0.63 |
| 75 | | | 0.80 | $10^{-4}$ |
| 250 | | | | 0.21 |

**Figure 6B**. 1D CMS velocity ($n_{motor}$ = 1000)

| $n_{clutch}$ | 25 | 75 | 250 | 750 |
|---|---|---|---|---|
| 8 | 0.05 | $10^{-8}$ | 0.63 | 0.56 |
| 25 | | 0.37 | 0.99 | 0.007 |
| 75 | | | 0.39 | $10^{-5}$ |
| 250 | | | | 0.09 |

**Figure 6C**. cRGD experiment motility

| [cRGD] | 0.1 | 0.3 | 1 |
|---|---|---|---|
| 0 | 0.54 | 0.97 | 0.67 |
| 0.1 | | 0.99 | 0.09 |
| 0.3 | | | 0.41 |
| 1 | | | |

**Figure 6D**. cRGD experiment velocity

| [cRGD] | 0.1 | 0.3 | 1 |
|---|---|---|---|
| 0 | 0.0012 | 0.00088 | $10^{-14}$ |
| 0.1 | | 0.99 | $<10^{-15}$ |
| 0.3 | | | $<10^{-15}$ |
| 1 | | | |

**Figure 6E.** 1D CMS motility ($n_{clutch}$ = 750)

| $n_{motor}$ | 500 | 750 |
|---|---|---|
| 100 | 0.99 | $10^{-4}$ |
| 500 | | 0.0028 |
| 750 | | |

**Figure 6F**. 1D CMS velocity ($n_{clutch}$ = 750)

| $n_{motor}$ | 500 | 750 |
|---|---|---|
| 100 | 0.099 | 0.0079 |
| 500 | | 0.88 |
| 750 | | |

Pairwise obtained by Kruskal-Wallis test with Dunn-Sidǎk multiple comparisons correction.

Shading indicates statistical significance for a particular pairwise comparison.

### Supporting Figures

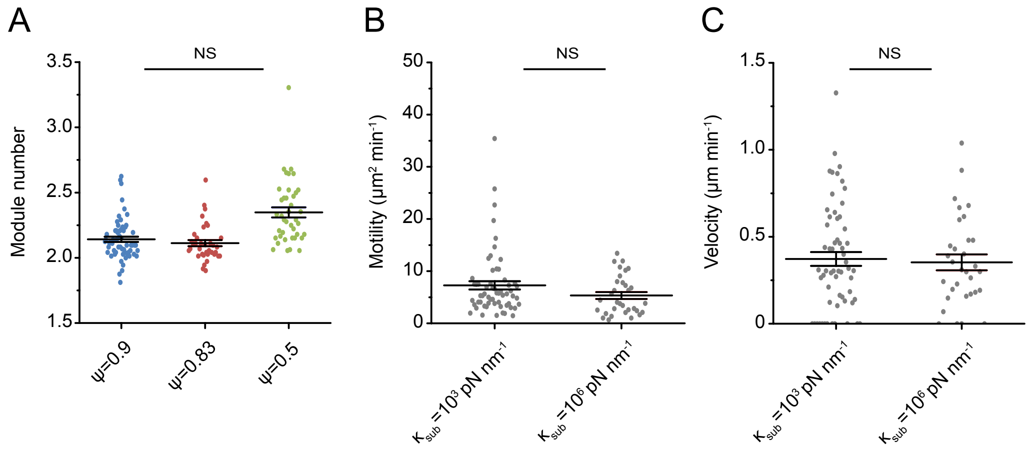


**Figure S1.** Additional 1D CMS outputs and effects of variable substrate stiffness on migration behaviors. Related to **Figures 1** and **3**.

(A) Number of modules for the simulations with ψ = 0.9, 0.83, or 0.5 from **Figure 1**.

(B) Motility coefficient from 1D CMS runs where $\kappa_{sub} = 10^3$ pN nm$^{-1}$ or $\kappa_{sub} = 10^6$ pN nm$^{-1}$; n = 60, 24 simulations.

(C) Velocity for the individual simulations in panel A. All motility coefficients and velocities are measured from individual fits to a diffusion-convection model (Eqn. 10). Simulations with $\kappa_{sub} = 10^3$ pN nm$^{-1}$ are reproduced from the reference dataset in **Figures 1** and **3**. Error bars represent mean ± SEM. Statistical comparisons in panel A were facilitated by Kruskal-Wallis analysis of variance with subsequent Dunn-Sidăk test and in panels B and C by Mann-Whitney U test. NS indicates no significant difference, defined as $p > 0.01$.

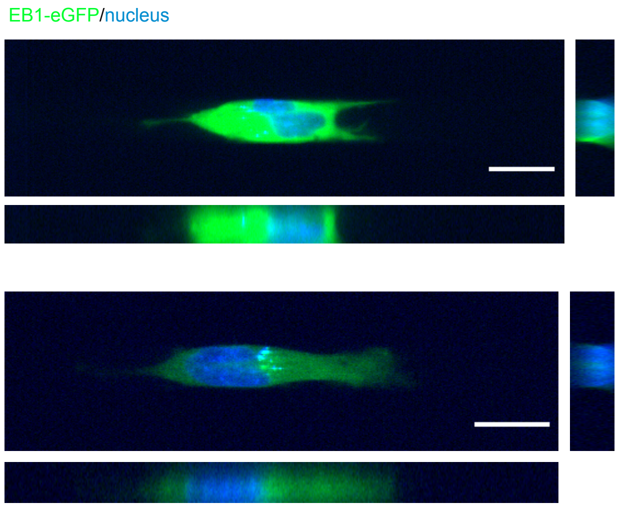


**Figure S2.** Confocal images of U251 human glioma cells in microchannels. Related to **Figure 2**.

Example confocal z-stacks of U251 cells expressing EB1-eGFP (green) and nucleus stain (blue). Images were acquired at 40x magnification. Images were oriented such that the channel inlets are to the left, while the outlets are to the right and adjusted for brightness/contrast in both green and blue channels. A view of the x-y plane is shown left, and an x-y slice through the center is shown at right. Total z-stack height, 11.43 μm; horizontal scale, 20 μm.

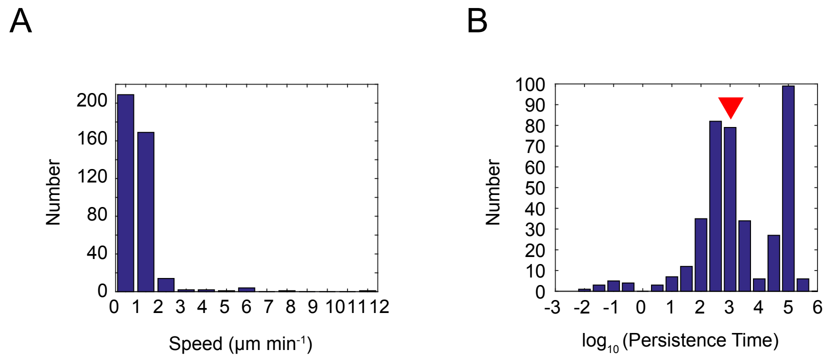


**Figure S3.** Persistent random walk fits yield persistence times that are longer than the typical experimental duration. Related to **Figures 2** and **3.**

(A) Individual cell speeds for the control dataset in **Figure 3**. Mean speed for this group: S = 0.74 ± 0.05 µm $min^{-1}$ or S = 12.3 ± 0.8 nm $s^{-1}$.

(B) Persistence time ($log_{10}$) for the fits in panel A. Red arrow indicates 1080 minutes (18 hrs), the maximum experimental duration. Mean $log_{10}$-transformed persistence time from individual fits: 3.1±0.25, corresponding to 1258 minutes.

### Supporting Movies

**Movie S1.** U251 glioma cells with fluorescently labeled nuclei migrating in microchannel devices.

Time-lapse images were collected every 5 minutes at 20x magnification with 2x2 binning (645 nm spatial sampling). Images were acquired in both the transmitted channel using phase contrast optics and LED fluorescence excitation (395 nm) using a DAPI/FITC/TxRed filter set. Scale bar, 50 µm.

**Movie S2**. U251 glioma cells expressing EGFP-actin and treated with vehicle control or latrunculin A migrating in microchannel devices.

Time-lapse images were collected every 5 minutes at 20x magnification with 2x2 binning (645 nm spatial sampling). Images were acquired in both the transmitted channel using phase contrast optics, plus LED fluorescence excitation (395 nm and 470 nm) using a DAPI/FITC/TxRed filter set. Conditions include DMSO vehicle (top), 50 nM latrunculin A (middle), and 500 nM latrunculin A (bottom). Scale bar, 50 µm.
